# Can a flux-based mechanism explain positioning of protein clusters in a three-dimensional cell geometry?

**DOI:** 10.1101/496364

**Authors:** Matthias Kober, Silke Bergeler, Erwin Frey

## Abstract

The plane of bacterial cell division must be precisely positioned. In the bacterium *Myxococcus xanthus*, the proteins PomX and PomY form a large cluster, which is tethered to the nucleoid by the ATPase PomZ and moves in a stochastic, but biased manner towards midcell, where it initiates cell division. Previously, a positioning mechanism based on the fluxes of PomZ on the nucleoid was proposed. However, the cluster dynamics was analyzed in a reduced, one-dimensional geometry. Here we introduce a mathematical model that accounts for the three-dimensional shape of the nucleoid, such that nucleoid-bound PomZ dimers can diffuse past the cluster without interacting with it. Using stochastic simulations, we find that the cluster still moves to and localizes at midcell. Redistribution of PomZ by diffusion in the cytosol is essential for this cluster dynamics. Our mechanism also positions two clusters equidistantly on the nucleoid. We conclude that a flux-based mechanism allows for cluster positioning in a biologically realistic three-dimensional cell geometry.

## I. INTRODUCTION

In bacteria, intracellular positioning of proteins is important for vital biological processes, including localization of the cell division machinery to midcell, as well as chromosome and plasmid segregation. The positioning systems responsible often involve P-loop ATPases such as ParA and MinD [1, 2]. These ATPases switch between an ATP- and ADP-bound state, which alters their subcellular localization: the ATP-bound form typically binds as a dimer to the nucleoid or membrane, while the ADP-bound form diffuses freely in the cytosol. Activating proteins stimulate the ATPase activity of the ParA-like proteins, which results in detachment of the protein from its respective scaffold in the ADP-bound form. Intracellular patterns of these ParA-like ATPases depend on the binding properties of the ADP- and ATP-bound forms, the localization of the stimulating proteins, and the cell geometry [3].

Recently, a protein system that includes a ParA-like ATPase has been identified, which regulates the localization of the cell division site in the bacterium *Myxococcus xanthus* [4–6]. The Pom system consists of three proteins: PomX, PomY and PomZ. The ATP-bound ParA-like ATPase PomZ binds as a dimer non-specifically to DNA. PomX and PomY form a macromolecular cluster, the PomXY cluster, which is tethered to the nucleoid via PomZ dimers. This tethering is transient, since PomX, PomY and DNA synergistically stimulate the ATPase activity of PomZ, which results in two ADP-bound PomZ monomers being released into the cytosol. Fluorescence labeling shows that, shortly after cell division, the PomXY cluster moves from a position close to one nucleoid end towards mid-nucleoid in a biased random walk-like movement that depends on PomZ. Once at mid-nucleoid, the Pom cluster positively regulates FtsZ ring formation [4].

Though the Pom system in *M. xanthus* and the Min system in *Escherichia coli* serve the same function (localization of the cell division plane), they differ in important respects: Dimeric MinD-ATP binds to the cell membrane, and MinC inhibits FtsZ ring formation outside of the midcell region, leading to negative regulation of mid-plane localization. In this regard, the Pom system is more similar to the Par system for low-copy-number plasmid and chromosome segregation, and related systems, e.g. those used to position chemotaxis protein clusters [7] and carboxysomes [8]. These systems utilize ParA-like ATPases that bind in the ATP-bound form to the nucleoid and regulate the movement of a cargo (e.g. protein cluster, partition complex or plasmids) to the required intracellular positions [9].

Previously, we introduced a one-dimensional mathematical model of the dynamics of the PomXY cluster in *M. xanthus* [4, 10] that takes the elasticity of the nucleoid [11, 12] into account. Based on this model, we proposed a mechanism that relies on PomZ dimer fluxes on the nucleoid to generate the experimentally observed cluster dynamics, which is midcell localization. A flux-based mechanism was originally proposed by Ietswaart et al. [13] for Par-mediated plasmid positioning. They showed that diffusive ParA fluxes can in principle explain equidistant spacing of plasmids along the nucleoid, because only when this is the case will the fluxes from either side of each plasmid balance up. Hence, macro-molecular objects can be evenly distributed on the nucleoid if they move in the direction from which the larger number of ParA proteins impinges upon them [13]. In our model of the Pom system, an analogous mechanism can localize the PomXY cluster to mid-nucleoid [4, 10].

It remained unclear, however, whether such a flux-based mechanism can also localize midcell if the full three-dimensional geometry of the nucleoid is accounted for (see Figure 1A). The reason is that, in contrast to the one-dimensional case, PomZ dimers can now more easily pass the cluster by without interacting with it. As a result, an asymmetry in the PomZ fluxes might be balanced. Here we show that the Pom cluster still localizes at midcell by a flux-based mechanism even if PomZ dimers can diffuse past the cluster.

**Figure 1.**
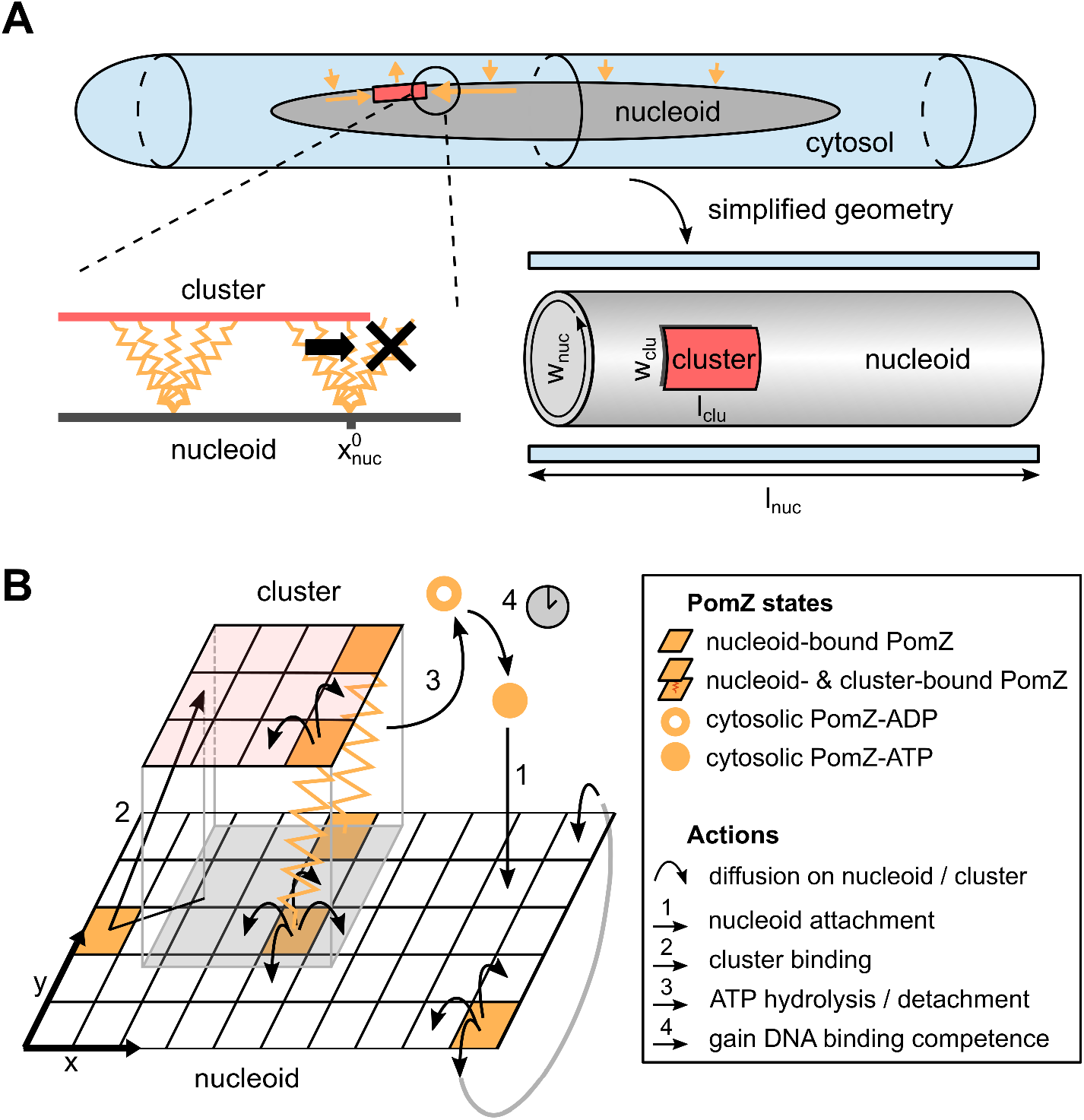
Mathematical model for PomXY cluster positioning, which accounts for the three-dimensional geometry of the cell. (A) Schematic representation of the geometry used in our model. Top: Sketch of a *M. xanthus* cell. Cytosolic, ATP-bound PomZ can bind to the nucleoid (orange arrows towards the nucleoid), and diffuses on the nucleoid. When bound to the PomXY cluster, PomZ hydrolyzes ATP and is released into the cytosol (orange arrow away from the nucleoid). These dynamics lead to a net diffusive flux of PomZ on the nucleoid towards the cluster, which is larger from the side with the larger cluster-to-nucleoid end distance (horizontal orange arrows) [13]. Bottom left: PomZ dimers, modeled as springs, can exert forces on the cluster by binding to the cluster in a deflected configuration and by encountering the cluster’s edge. For a particular nucleoid binding-site position, 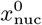, the positions available to PomZ’s cluster-binding site are limited if the dimer is located close to the cluster’s edge (black cross). This asymmetry can result in a force exerted on the cluster (black arrow). Bottom right: The model geometry derived from the biologically realistic three-dimensional cell. The nucleoid is modeled as an open cylinder and the cluster as an object of fixed size of rectangular shape. The cytosolic PomZ-ATP distribution is either assumed to be homogeneous or included effectively by modeling the cytosolic PomZ distribution along the long cell axis. (B) Schematic of the interactions of PomZ with the nucleoid and cluster considered in our model (see main text and SI text for details).

## II. RESULTS

### A. Flux-based mechanism for midcell localization

The dynamics of the PomXY cluster on the nucleoid crucially depends on the PomZ dynamics, as the cluster is tethered to the nucleoid via PomZ dimers [4]. Based on the biochemical processes suggested by experiments [4], we model the dynamics of PomZ as follows (see Figure 1B and SI text for details). ATP-bound PomZ dimers bind to the nucleoid with rate *k*_on_ (action 1 in Figure 1B). Once on the nucleoid, they diffuse with diffusion constant *D*_nuc_. The PomZ dimers are modeled effectively as springs to account for the elasticity of the chromosome and the Pom proteins (as in [12, 14]). For simplicity we will refer to the PomZ dimers as springs in the following, although the elasticity mainly originates from the nucleoid.

A nucleoid-bound PomZ dimer can attach to the cluster with rate *k_a_* either ‘orthogonally’, or obliquely in an extended configuration (action 2 in Figure 1B). We assume that cluster-bound PomZ can diffuse on both the nucleoid and the PomXY cluster. However, the freedom of movement of nucleoid- and cluster-bound PomZ is restricted due to the energy cost involved in stretching the spring (for details see SI text). Cluster-bound PomZ dimers remain attached to the cluster until they are released into the cytosol upon ATP hydrolysis, which is stimulated by PomX, PomY and DNA. ATP hydrolysis leads to a conformational change in the PomZ dimer and triggers the release of two ADP-bound PomZ monomers into the cytosol [4]. In our model, we combine these processes into one by using a single rate *k_h_* to describe the detachment of cluster-bound PomZ-ATP dimers into the cytosol as monomers (action 3 in Figure 1B). Before PomZ can rebind to the nucleoid, it must first bind ATP and dimerize. This introduces a delay between detachment from and reattachment to the nucleoid, which allows for spatial redistribution of the quickly diffusing cytosolic PomZ dimers in the cell (action 4 in Figure 1B).

The cluster dynamics, which we approximate as over damped, is determined by the forces exerted by the PomZ dimers on the cluster and the friction coefficient of the cluster (see SI text). Previously we suggested a flux-based mechanism for its midcell positioning in a onedimensional model geometry [4, 10], which can be summarized as follows. The PomZ dimers, which are modeled as springs, can exert net forces on the cluster in two different ways. They can attach to the cluster in a stretched configuration and thereby exert forces on the cluster (similar to the DNA-relay mechanism proposed for Par systems, [12]). However, for PomZ dimers that quickly diffuse on the nucleoid (as observed experimentally in *M. xanthus* cells) the initial deflection of the spring relaxes within a short time, such that only a small net force is exerted on the cluster. In this case, forces are mainly generated by cluster-bound PomZ dimers that encounter the cluster’s edge (Figure 1A, bottom left) [10]. Every time a nucleoid- and cluster-bound PomZ dimer reaches the cluster’s edge, the nucleoid binding site can diffuse beyond the cluster region, whereas the cluster binding site is restricted in its movement, which on average results in a net force on the cluster. These forces are such that a protein that reaches the cluster from the right exerts a force, which ‘drags’ the cluster to the right and vice versa.

The fact that PomZ dimers can exert forces on the cluster is not enough to explain the movement of the cluster towards midcell. Here, the asymmetry in the PomZ dimer density on the nucleoid, which prevails as long as the cluster is located off center, i.e. not already at mid-nucleoid, is crucial. Since PomZ dimers can attach anywhere on the nucleoid but can detach only when they make contact with the cluster (as stimulation of their ATPase activity depends on PomX, PomY and DNA), a non-equilibrium flux of PomZ dimers in the system can be maintained. In particular, this flux includes a diffusive flux of PomZ dimers along the nucleoid towards the cluster. For a cluster located off-center the diffusive PomZ fluxes into the cluster from either side will differ [13]. Since the average forces applied by PomZ dimers arriving from the right and left sides of the cluster act in opposite directions, the difference in the PomZ fluxes from the two sides determines the net force exerted on the cluster [10]. Taken together, these factors combine to drive a self-regulated midcell localization process as long as the PomZ dynamics is fast compared to the cluster dynamics and, if this is not the case, lead to oscillatory cluster movements along the nucleoid [10].

### B. A three-dimensional model for midcell localization

To understand how the geometry of the nucleoid and the size of the PomXY cluster affect the cluster dynamics and thereby test whether a flux-based mechanism is feasible in a biologically realistic three-dimensional geometry, we investigated the mathematical model illustrated in Figure 1. Here, the nucleoid and the PomXY cluster are approximated as a cylindrical object and a rectangular sheet, respectively. Since experiments in *M. xanthus* cells suggest that the cluster is large [4] we assume that the cluster, tethered to the nucleoid via PomZ dimers, moves over the nucleoid’s surface and does not penetrate the bulk of the nucleoid. Moreover, we assume that PomZ dimers also bind to and diffuse on the nucleoid’s surface only.

The cylindrical geometry of the nucleoid is mathematically implemented by a rectangular sheet with periodic boundary conditions for the PomZ movements along the short cell axis (*y* direction) and reflecting boundary conditions along the long cell axis (*x* direction, Figure 1B). The cluster is modeled as a rectangle with reflecting boundaries at its edges for the PomZ dimer movement. We refer to the extension of the cluster along the long and short cell axis as the cluster’s length, *l*_clu_, and width, *w*_clu_, respectively (see Figure 1A).

Besides the nucleoid and the cluster, the cytosol needs to be accounted for in our model, as PomZ dimers cycle between a nucleoid-bound and cytosolic state. We expect the cytosolic diffusion constant of PomZ to be of the same order as that of MinD proteins in *E. coli* cells, *D*_cyt_ ≈ 10 μm^2^s^-1^ [15]. For ParA ATPases involved in chromosome and plasmid segregation, a delay between the release of ParA from the nucleoid into the cytosol and the re-acquisition of its capacity for non-specific DNA binding is observed experimentally [16]. Upon ATP hydrolysis and release of two ADP-bound ParA monomers into the cytosol, ParA must bind ATP, dimerize and then regain the competence to bind non-specifically to DNA before it can reattach to the nucleoid. The last step, a conformational change in the ATP-bound ParA dimer, is the slowest of these processes, occurring on a time scale of the order of minutes, *τ* ≈ 5min [16]. Since PomZ is ParA-like, we expect the PomZ ATPase cycle to be similar to the ParA cycle, and therefore assume the corresponding reaction rates to be of the same order of magnitude. This yields the following estimate for the diffusive length of cytosolic PomZ, 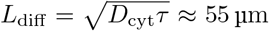, which is significantly larger than the average cell length of a *M. xanthus* cell of 7.7 μm [4]. Hence, the assumption of a well-mixed PomZ density in the cytosol is justified [3].

However, it is not known how the cytosolic distribution of PomZ affects the cluster dynamics in our proposed flux-based mechanism. In particular, does a realistic, non-uniform distribution increase or reduce the speed of the cluster movement towards midcell? To obtain a qualitative answer to this question, we accounted for the cytosolic PomZ distribution in a simplified way by focusing on the variation in PomZ density along the long cell axis and approximating the density along the short cell axis as uniform (Figure 1A, bottom right). In section II E we discuss how the cluster dynamics is affected when PomZ’s diffusion constant in the cytosol is reduced (which leads to deviations from the uniform PomZ distribution), but for now we assume a homogeneous PomZ-ATP distribution in the cytosol.

### C. A flux-based mechanism can explain midcell positioning in three dimensions

Our simulations using the three-dimensional model geometry show that the net force exerted by the PomZ dimers on a stationary cluster (i.e. a cluster with fixed position) is still directed towards mid-nucleoid (Figure 2A). The asymmetry in the forces can be attributed to an asymmetry in the PomZ fluxes along the nucleoid into the cluster. This finding indicates that the cluster movement is biased towards midcell in the three-dimensional model geometry also. Hence, a flux-based mechanism is also conceivable in three dimensions. Indeed, the simulated PomXY cluster trajectories show movement towards and localization at mid-nucleoid (Figure 2B). If the cluster’s width does not cover the complete nucleoid circumference, the movement towards midcell takes longer than in the one-dimensional case (Figure 2B). The forces along the short cell axis direction balance (Figure 2A) and hence no bias in the cluster movement along the short axis is expected for a slowly moving cluster and fast PomZ dynamics. The simulated cluster trajectories show diffusive motion along the short cell axis direction for the parameters given in Table S1 (see SI text).

**Figure 2.**
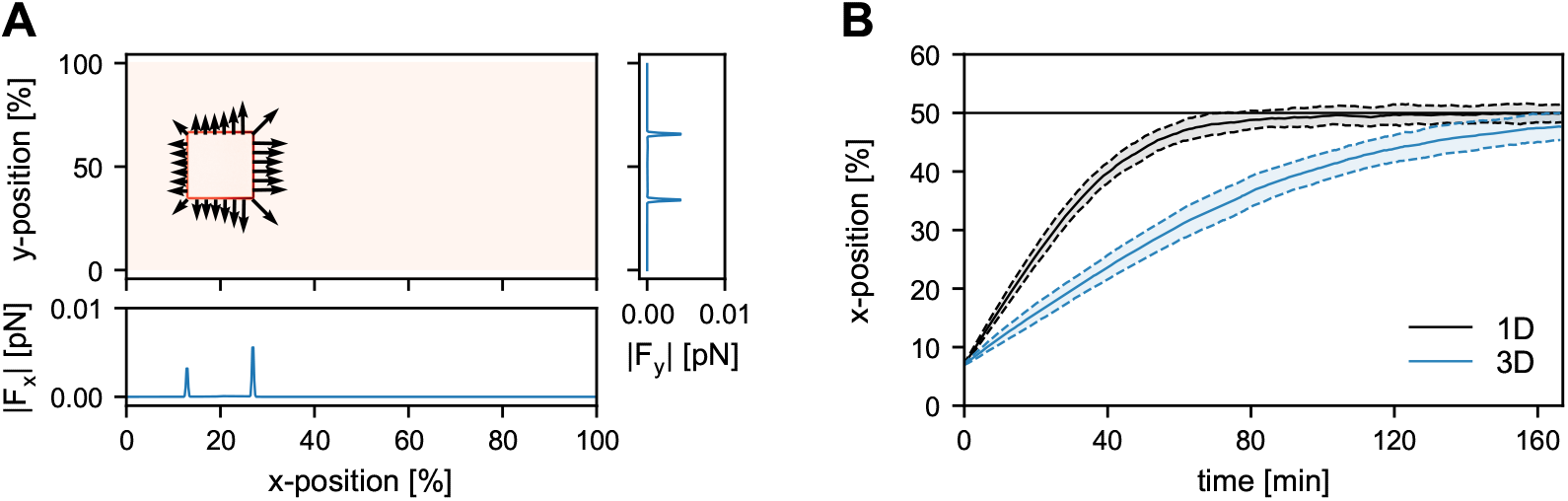
A flux-based mechanism can explain midcell localization in a three-dimensional cell geometry. (A) For a fixed cluster position (here at 20% nucleoid length), the average forces exerted by PomZ dimers on the cluster are plotted per nucleoid lattice site. The color code shows the magnitude of the average force vector, which is highest at the cluster’s edges (darker red indicates higher values). At these edges, the average force vectors are plotted, 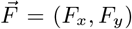. The average *x*- and *y*-component of the force, *F_x_* and *F_y_*, (summed over all *y*- and *x*-positions, respectively), are shown in the lower and right panels, respectively. (B) Comparison of the average cluster trajectories along the cell’s long axis, obtained from the three-dimensional model and its one-dimensional counterpart studied previously [10] (see also Figure S3). We averaged over an ensemble of 100 simulations. The shaded regions depict one standard deviation around the mean density value. In all cases, the cluster is initially positioned at the left edge of the nucleoid such that it overlaps entirely with the nucleoid (at 7% of the nucleoid length). Mid-nucleoid is indicated by the horizontal black line. For the dynamics of the cluster in a single simulation see Movie S1. The parameter values used in the simulations are given in Table S1.

### D. The cluster’s linear dimensions determine the time taken to reach midcell

In this section we ask how the arrival time of the cluster at mid-nucleoid depends on the linear dimensions of the cluster. Our simulations show that the larger the cluster’s length or width, the faster the cluster moves towards midcell (Figure 3A). In the following we discuss how these two observations can be explained heuristi-cally.

**Figure 3.**
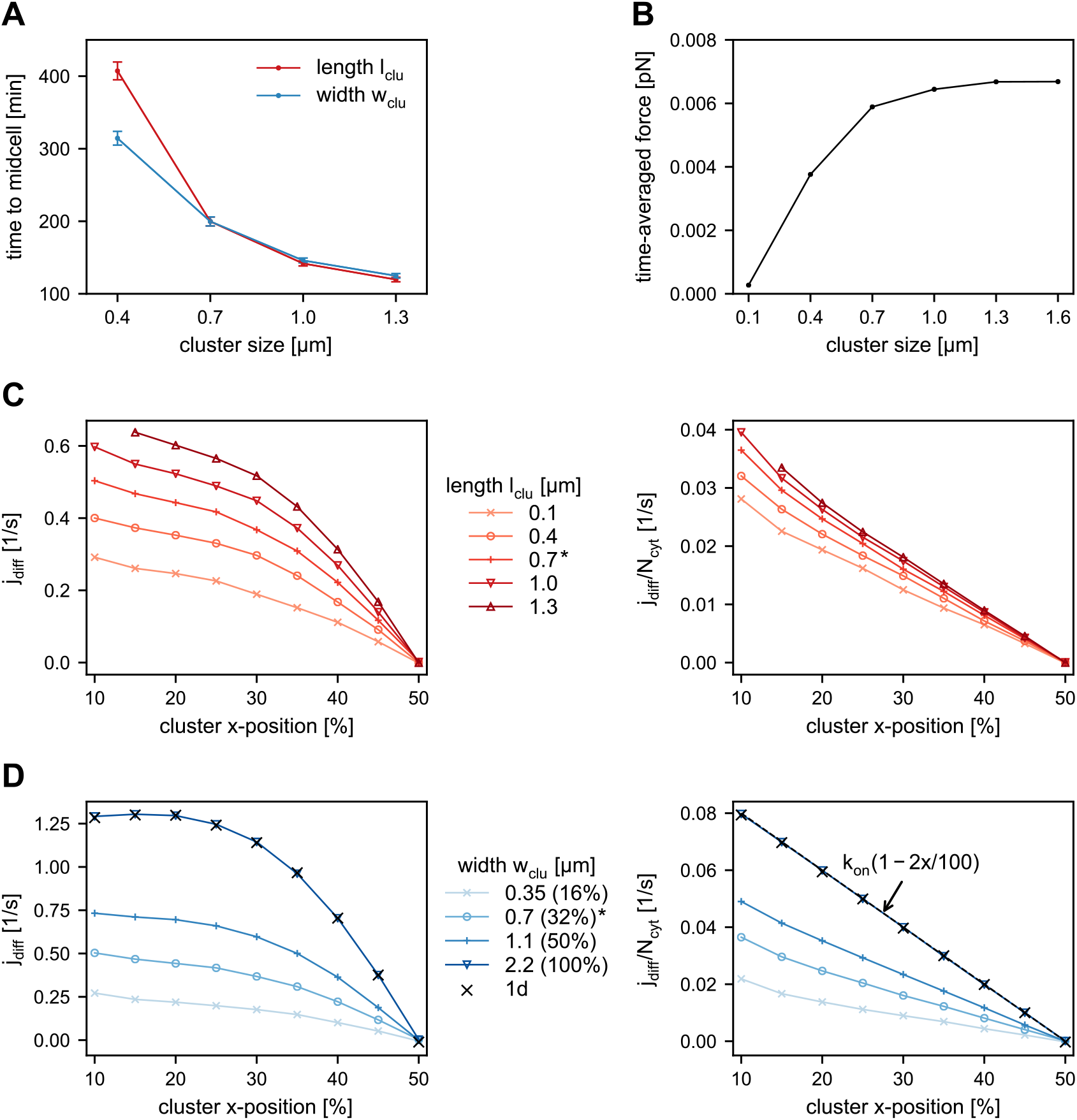
Dependence of the time needed to reach midcell on the size of the cluster. (A) Average first passage time of the PomXY cluster to reach mid-nucleoid for different cluster sizes. In all simulations the cluster starts at 13% of nucleoid length, which corresponds to the leftmost position possible such that for all cluster sizes considered a full overlap with the nucleoid is ensured. The error bars show the standard error of the mean. (B) Ensemble average of the time-averaged force exerted by a single PomZ dimer on the cluster for a one-dimensional model geometry and different cluster lengths. (C)-(D) The PomZ flux difference into the cluster, *j*_diff_, along the long cell axis for a cluster at a fixed position, which is varied from 10% to 50% of nucleoid length, is shown (see also Figure S2). In (C), the cluster’s length is varied and in (D) the cluster’s width. For a ring-shaped cluster, *w*_clu_ = *w*_nuc_, the simulation results agree with those from the one-dimensional model (black crosses). To understand how the cluster’s length and width affect the flux difference, the values are scaled with the number of PomZ dimers in the cytosol, *N*_cyt_, for each cluster position separately (Figures on the right). In (D) an analytical estimate for the flux difference is plotted (dashed line), which agrees with the simulation results for a ring-shaped cluster (see SI text for details). If not given explicitly, the parameter values used are those listed in Table S1 (data marked with a star).

The net force applied to the cluster by PomZ dimers depends on the average force exerted by a single PomZ dimer and on the flux difference in PomZ dimers arriving at the cluster [10]. The latter can be regarded as the frequency of PomZ interactions with the cluster that lead to a net force contribution.

The flux difference into the cluster increases for larger or broader clusters for two reasons. First, increasing the length or width of the cluster results in an increased number of cluster-bound PomZ dimers, as the cluster becomes more accessible to nucleoid-bound PomZ dimers. This increase in cluster-bound PomZ dimers leads to a larger turnover of PomZ dimers cycling between the nucleoid-bound and cytosolic state, which also increases the flux difference. Second, the larger and wider the cluster, the less likely it is that PomZ dimers will diffuse past it without attaching, which would otherwise reduce the flux difference into the cluster along the long cell axis.

The forces exerted by PomZ dimers also depend on the linear dimensions of the cluster. Increasing the cluster size increases the average force exerted by a single PomZ dimer, until a maximal force is reached (Figure 3B). This dependence can be explained by diffusion of PomZ dimers bound to the cluster: The smaller the cluster size, the more likely it is that a PomZ dimer attaching close to one edge of the cluster will reach the other edge before detaching into the cytosol. Since the PomZ dimer then exerts forces in both directions along the long cell axis, the time-averaged force it applies to the cluster is reduced.

Based on these observations, we can rationalize the dependence of the arrival time at mid-nucleoid on the length of the cluster as follows. An increase in cluster length (while keeping its width constant) increases the numbers of PomZ dimers interacting with the cluster, and hence the flux difference along the long cell axis. Indeed, our simulation results show that the longer the cluster, the larger the difference in the PomZ dimer fluxes from either side along the direction of the long cell axis (Figure 3C). To test whether the increased flux of PomZ dimers in the system is the main determinant for the observed increase in the flux difference, we scaled the fluxes by the number of PomZ dimers in the cytosol, *N*_cyt_, which is proportional to the flux of PomZ dimers onto the nucleoid. Upon rescaling, the flux differences decay approximately linearly with the cluster position (Figure 3C, right). However, longer clusters still show the largest flux differences. This phenomenon can be attributed to the fact that for shorter clusters PomZ dimers are more likely to diffuse past the cluster.

For a longer cluster not only the diffusive flux of PomZ dimers into the cluster, but also the force exerted by a single PomZ dimer on the cluster is increased (Figure 3B). Hence, frequency and magnitude of forces exerted on the cluster are increased, implying a larger net force, which explains the shorter arrival times of the cluster at mid-nucleoid (Figure 3A).

Next, we discuss the dependence of the arrival time on the cluster width. As in the case of an increase in length, the overall turnover of PomZ dimers increases with cluster width. The width of the cluster determines how many PomZ dimers approach and attach to the cluster from the directions of the long and the short cell axes, respectively, and how many pass the cluster without interacting with it. The broader the cluster, the smaller the flux of PomZ dimers that can diffuse past the cluster without attaching. Our simulation results show an increased PomZ flux difference when the cluster width is increased from 16% to 100% of the nucleoid’s circumference (Figure 3D). When the cluster covers the entire circumference of the nucleoid, we call it a ring. Rescaling of the flux differences with the number of PomZ dimers in the cytosol, *N*_cyt_, leads to values that are still larger for wider clusters, due to the reduced flux of PomZ dimers past the cluster (Figure 3D, right). The forces exerted by single PomZ dimers in the direction of the long cell axis direction are not affected by a change in the cluster width. Hence, the decrease in arrival time for wider clusters can be explained by the increased PomZ flux difference into the cluster along the long cell axis alone.

### E. Cytosolic diffusion ensures fast midcell positioning of the cluster

So far, we have assumed that the cytosolic PomZ distribution is spatially uniform. Now we investigate how the cluster dynamics change when spatial heterogeneity in the cytosolic PomZ distribution is included in the model. To this end, we explicitly incorporate the cytosol by approximating the cytosolic volume as a one-dimensional layer of the same length as the nucleoid (see Figure 1A) and formulate reaction-diffusion equations for the density of cytosolic PomZ-ADP (*c_D_*) and PomZ-ATP (*c_T_*) along the long cell axis. For simplicity, we consider only an active (ATP-bound) and inactive (ADP-bound) conformation of PomZ, and disregard any explicit monomeric and dimeric states of PomZ in the cytosol (for details see SI text). The stationary solution for *c_T_*(*x*; *x_c_*) deviates most from a uniform distribution, the smaller the cytosolic diffusion constant (Figure 4A). In the limit of infinitely large cytosolic PomZ diffusion constants, the cytosolic PomZ distribution becomes spatially uniform.

**Figure 4.**
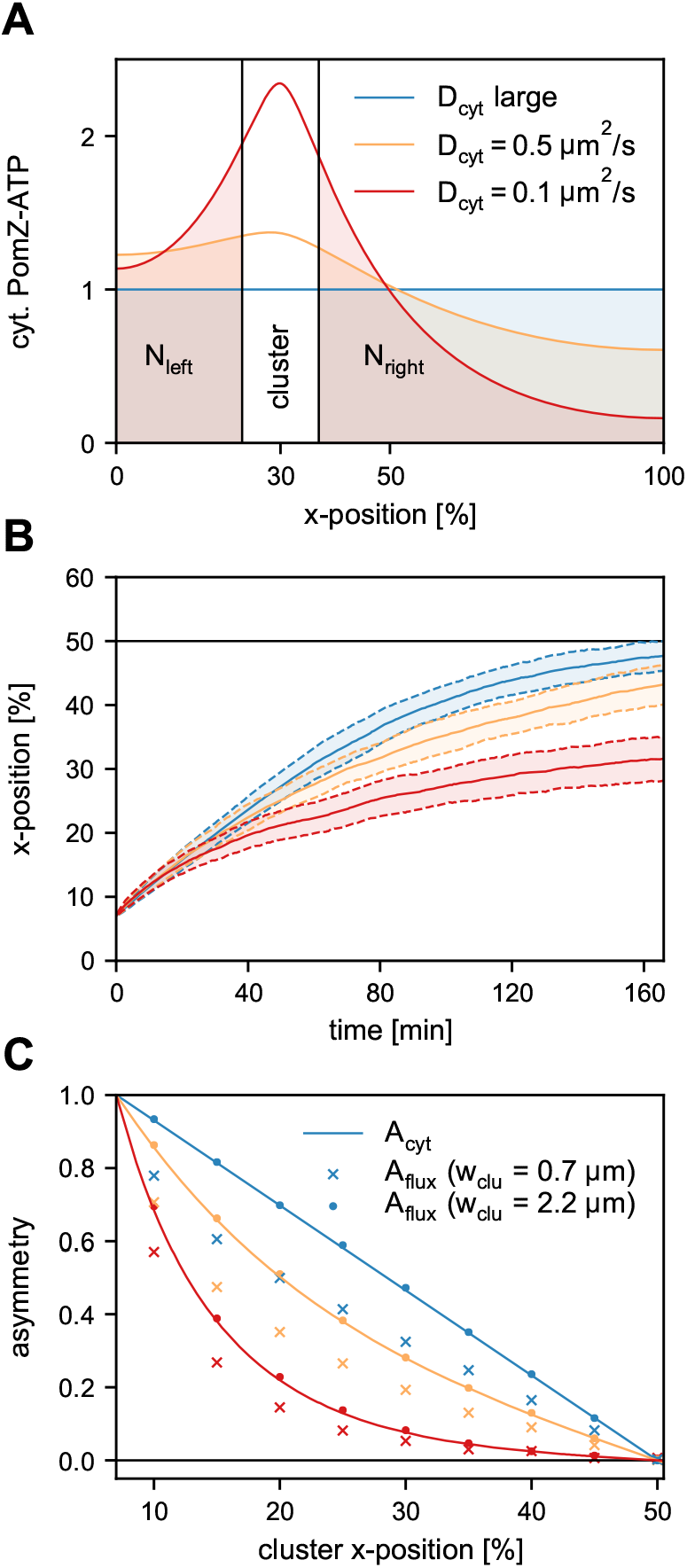
Fast cytosolic diffusion accelerates midcell localization. (A) Cytosolic PomZ-ATP distribution along the long axis (*x* axis) for different cytosolic PomZ diffusion constants *D*_cyt_ (see also Figure S1). Integrating the distributions along the long cell axis left and right of the cluster yields the total number of PomZ-ATP proteins left and right of the cluster, *N*_left_ and *N*_right_. (B) Average cluster trajectories along the *x* direction for the different cytosolic PomZ diffusion constants in (A). The shading denotes the regions of one standard deviation around the average trajectories. Mid-nucleoid is indicated by the solid black line. In the simulations, the clusters are positioned initially such that the left edge of the cluster coincides with the left edge of the nucleoid. (C) Asymmetry measure of the number of cytosolic PomZ-ATP left and right of the cluster, *A*_cyt_ (Equation 1) (solid lines), compared to the PomZ flux asymmetry into the cluster, *A*_flux_ (Equation 4), for different cytosolic PomZ diffusion constants. The dots indicate the flux asymmetry into a cluster that forms a ring. Crosses indicate the asymmetry for a cluster with a width of 32% of the nucleoid’s circumference (same value as in Table S1). We averaged over 100 runs of the simulation. The parameter values given in Table S1 are used if not explicitly stated otherwise.

To investigate the effect of the cytosolic PomZ distribution on the cluster’s trajectory, we replaced the spatially uniform cytosolic PomZ-ATP distribution by *c_T_*(*x*; *x_c_*), in our three-dimensional model. Since *M. xanthus* cells are rod-shaped with the length being much larger than the width, we approximate the cytosolic PomZ-ATP distribution along the short cell axis as uniform. With this spatially heterogeneous attachment rate to the nucleoid, our simulations show that for a larger deviation of the cytosolic PomZ distribution from a spatially uniform one (decreasing *D*_cyt_), the movement of the clusters is less biased towards mid-nucleoid (Figure 4B). We conclude that the cytosolic PomZ distribution has an impact on the cluster trajectories and the velocity of the cluster towards mid-nucleoid is maximal for a uniform cytosolic PomZ-ATP distribution.

How does the cytosolic PomZ distribution affect the cluster’s movement? The flux of PomZ dimers onto the nucleoid region to the left of the cluster and the diffusive flux of PomZ dimers into the cluster from the left are equal for a stationary cluster in the one-dimensional model geometry, and vice versa for the right side (see SI text). Since the total flux of PomZ dimers onto the nucleoid to the left and right of the cluster depends on the cytosolic PomZ-ATP distribution, we expect the diffusive PomZ fluxes into the cluster to depend on this distribution as well. To investigate this further, we define the following asymmetry quantity

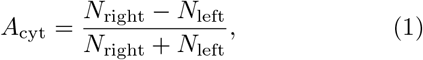

which compares the total numbers of cytosolic PomZ-ATP left and right of the cluster (see Figure 4A):

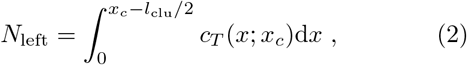

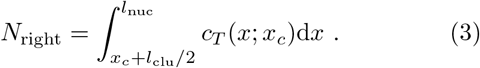

The corresponding asymmetry in the PomZ fluxes into the cluster from the left, *j*_left_, and right, *j*_right_, sides is given by

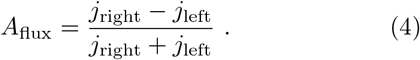

We measured this flux asymmetry in our simulations for two scenarios. First, for a PomXY cluster that forms a ring around the nucleoid and second, for a cluster that does not entirely encompass the nucleoid’s circumference. We find that the asymmetry in the flux, *A*_flux_, obtained from simulations for a cluster that forms a ring, and the corresponding asymmetry in the cytosolic PomZ-ATP density, *A*_cyt_, agree nicely (Figure 4C). Hence, an asymmetry in the cytosolic distribution of PomZ-ATP is directly reflected in the diffusive PomZ fluxes into the cluster. For an infinitely large cytosolic PomZ diffusion constant, both asymmetry measures decay linearly when the cluster position is varied from far off-center towards mid-nucleoid. Decreasing the cytosolic PomZ diffusion constant results in asymmetry curves that decay faster than linearly towards zero (Figure 4C). For a cluster that does not cover the whole nucleoid’s circumference, the asymmetry in the PomZ fluxes into the cluster is smaller than the asymmetry in the cytosolic PomZ-ATP concentration (Figure 4C). This can be attributed to the reduction in the diffusive fluxes of PomZ dimers into the cluster along the long cell axis, as discussed before. The reduced asymmetry in the diffusive PomZ fluxes explains the less biased movement of the cluster towards mid-nucleoid for smaller cytosolic PomZ diffusion constants (Figure 4B).

### F. Two clusters localize at one- and three-quarter positions

Motivated by equidistant positioning of multiple cargoes, such as plasmids [13, 17–20], we also considered the dynamics of two PomXY clusters in the threedimensional model geometry. Our simulation results show equidistant positioning of the two clusters (see SI text, Figure S4 and Movie S2).

## III. DISCUSSION

In this work we investigated a mathematical model for midcell localization in *M. xanthus* using a biologically realistic three-dimensional geometry for the nucleoid. Whether or not a flux-based mechanism can position macromolecular objects when the ATPases (here PomZ) can diffuse past a cargo (here PomXY cluster) has been questioned, because the fluxes into the cargo might equalize [13]. We showed that if PomZ dimers can diffuse past the PomXY cluster, there is still a flux difference into the cluster (for a cluster positioned off-center), which leads to a bias in the cluster movement towards mid-nucleoid. Hence, we conclude that a flux-based mechanism can explain midcell positioning of one or equidistant localization of several Pom clusters even if PomZ can diffuse past the cluster.

To investigate the effect of the flux of PomZ dimers past the cluster, we studied the dependence of cluster dynamics on its width and length. We find that increasing the cluster length or width shortens the time taken for the cluster to reach mid-nucleoid. This can be attributed to an overall increase in the flux difference, reduced flux past the cluster and the larger forces single PomZ dimers exert, on average, on larger and wider clusters.

Our simulation data further demonstrates that fast cytosolic diffusion of PomZ proteins reduces the time taken to find midcell. This finding is in agreement with previous results for dynamic protein clusters in bacterial cells [21]. The asymmetry in the cytosolic PomZ density left and right of the cluster along the long cell axis is reflected in the diffusive flux difference of PomZ dimers into the cluster, and thus influences the cluster dynamics. If PomZ diffuses quickly in the cytosol, the cytosolic PomZ-ATP distribution becomes spatially uniform. In this case, the flux of PomZ dimers onto the nucleoid scales with the length of the nucleoid regions left and right of the cluster. This results in the largest PomZ flux differences (for an off-center cluster) compared to spatially non-uniform cytosolic PomZ-ATP distributions. Interestingly, spatial redistribution of proteins in the cytosol is also found to be important for other pattern-forming systems, including Min protein pattern formation [22, 23] and ParA-mediated cargo movement [16].

Le Gall et al. [24] have shown that partition complexes as well as plasmids move within the nucleoid volume. In contrast, based on the large size of the PomXY cluster, we assumed that the movement of the cluster, tethered to the nucleoid via PomZ dimers, is restricted to the surface of the nucleoid. To verify our assumption, the position of the cluster relative to the nucleoid needs to be measured *in vivo* using e.g. super-resolution microscopy. In addition, since PomZ dimers are much smaller than the PomXY cluster, they might be able to diffuse within the nucleoid volume even though the cluster does not. It would be interesting to investigate this aspect further.

In summary, we have shown that a flux-based mechanism can explain midcell localization of one, and equidistant positioning of two clusters in a model geometry that allows the ATPase PomZ to diffuse past the clusters on the nucleoid. This observation is also important for other positioning systems, such as the Par system, which equidistantly spaces low-copy-number plasmids along the nucleoid. Understanding the differences and similarities between these positioning systems will help us to understand the generic mechanisms underlying the localization patterns of cargoes inside the cell.

## Supporting information

## ACKNOWLEDGEMENTS

The authors thank Isabella Graf, Emanuel Reithmann, Christoph Brand, Dominik Schumacher and Lotte Søgaard-Andersen for helpful discussions. This research was supported by a DFG fellowship through the Graduate School of Quantitative Biosciences Munich, QBM (SB, EF), the Deutsche Forschungsgemeinschaft (DFG) via project P03 within the Transregio Collaborative Research Center (TRR 174) “Spatiotemporal Dynamics of Bacterial Cells” (SB, EF), and the German Excellence Initiative via the program “Nanosystems Initiative Munich” (EF).

