## Supplementary material for "Can a flux-based mechanism explain positioning of protein clusters in a three-dimensional cell geometry?"

### Supplemental Material

#### I. DETAILS ON THE MATHEMATICAL MODEL

##### 1. Attachment rate of PomZ to the nucleoid

In our simulations we either assume that the PomZ density in the cytosol is homogeneous such that the flux of PomZ dimers onto the nucleoid is constant along the nucleoid or we account for the cytosolic PomZ distribution in a simplified manner. In the former case, a cytosolic PomZ dimer attaches to each lattice site of the nucleoid, also where the cluster is located, with the same rate. This rate is given by the attachment rate to the entire surface of the nucleoid, $k_{\text{on}}$, divided by the number of lattice sites $l_{\text{nuc}} \cdot w_{\text{nuc}}/a^2$. By $a$ we denote the lattice spacing, which has the same value in $x$- and $y$-direction. In the latter case, when we account for the cytosolic PomZ distribution, we replace the homogeneous with the steady-state cytosolic PomZ-ATP distribution in our model (see section I 5). Here, we use a one-dimensional solution for the cytosolic PomZ-ATP density, which describes the variation of the protein density along the long cell axis. In our model, which has a three-dimensional geometry, we assume that the cytosolic PomZ-ATP density is uniform along the short cell axis and changes along the long cell axis according to the one-dimensional distribution.

##### 2. Attachment of PomZ to and detachment from the PomXY cluster

A nucleoid-bound PomZ dimer can bind to a lattice site on the PomXY cluster. Since PomZ dimers are modelled as springs to account for the elasticity of the nucleoid, they can attach to the cluster not only ‘orthogonally’, but also in a stretched configuration. We approximate the rate for a PomZ dimer, bound to the nucleoid at position $\vec{x}_{\text{nuc}}$, to attach to the cluster at position $\vec{x}_{\text{clu}}$, as follows:

$$k_a(\vec{x}_{\text{clu}}) = k_a^0 \cdot \exp\left[-\frac{1}{2}\,\beta k(\vec{x}_{\text{clu}} - \vec{x}_{\text{nuc}})^2\right] . \tag{S1}$$

The rate for attachment to a lattice site at position $\vec{x}_{\text{clu}}$ on the cluster is then given by $k_a(\vec{x}_{\text{clu}}) \cdot a^2$. In the formula above, we multiply the constant rate $k_a^0$ with a Boltzmann factor corresponding to the distribution of the elongation of a spring in a thermal heat bath with temperature $T$. Here, $k$ denotes the effective spring stiffness of the PomZ dimers and $\beta$ is the inverse of the thermal energy, $\beta = 1/k_B T$. The positions of the cluster and nucleoid binding sites of the PomZ dimer are denoted as $\vec{x}_{\text{clu}}$ and $\vec{x}_{\text{nuc}}$, respectively.

We expect that the position of a cluster, which is tethered to the nucleoid, does not change remarkably in the direction perpendicular to the nucleoid’s surface because the cluster is unlikely to penetrate into the nucleoid’s volume due to its large size and, on the other hand, cannot move far away from the nucleoid due to the tethering. Hence we neglect the forces that a PomZ dimer exerts on the cluster in the direction perpendicular to the nucleoid’s surface by approximating the positions of the cluster and nucleoid binding site by their projections on the rectangular sheet representing the nucleoid, i.e. $\vec{x}_{\text{clu}} = (x_{\text{clu}}, y_{\text{clu}})$, and $\vec{x}_{\text{nuc}} = (x_{\text{nuc}}, y_{\text{nuc}})$.

The rate for a PomZ dimer located at $\vec{x}_{\text{nuc}}$ to attach, with its second binding site, to any site of the cluster (“total attachment rate”) is then given by integration of $k_a(\vec{x}_{\text{clu}})$ over all possible cluster binding sites:

$$k_a^{\text{tot}} = \int_{\mathcal{A}_{\text{clu}}} k_a^0 \cdot \exp\left[-\frac{1}{2}\,\beta k(\vec{x}_{\text{clu}} - \vec{x}_{\text{nuc}})^2\right] \mathrm{d}\vec{x}_{\text{clu}} \approx \begin{cases} k_a^0 \cdot \frac{2\pi}{\beta k}, & \text{if } \vec{x}_{\text{nuc}} \in \mathcal{A}_{\text{clu}} \ , \\ 0, & \text{otherwise} \ . \end{cases} \tag{S2}$$

Here, $\mathcal{A}_{\text{clu}}$ denotes the area on the nucleoid that is covered by the cluster. Since the Boltzmann factor decays quickly ($1/\sqrt{\beta k} = 0.01\,\mu\text{m}$ for the spring stiffness used, see Table S1), we can neglect the boundaries of the region $\mathcal{A}_{\text{clu}}$ for $\vec{x}_{\text{nuc}} \in \mathcal{A}_{\text{clu}}$ and approximate the attachment rate of a PomZ dimer bound to the nucleoid outside of the cluster region by zero.

Due to the fast decay of the exponential factor with an increase in $|\vec{x}_{\text{clu}} - \vec{x}_{\text{nuc}}|$, we can also save computation time by introducing a cut-off distance above which we set the attachment rate to zero. The cut-off is defined as the smallest distance $\Delta x = |\vec{x}_{\text{clu}} - \vec{x}_{\text{nuc}}|$ for which the attachment rate per lattice site, $k_a \cdot a^2$, is smaller than $10^{-5}\,\text{s}^{-1}$. This value is chosen such that the average number of times this event occurs during the time the cluster takes to reach midcell ($\approx 4800\,\text{s}$) is 0.048, i.e. much lower than one.

PomZ dimers are captured at the cluster until they are released into the cytosol upon ATP hydrolysis. In our model, we combine the different processes (ATP hydrolysis, conformational change in PomZ and detachment of PomZ-ADP into the cytosol) into one effective detachment process and denote the corresponding rate as the ATP hydrolysis rate $k_h$, which we assume to be independent of the degree of stretching of the dimer.

##### 3. Hopping of PomZ dimers on the nucleoid and the PomXY cluster

In our model we assume that nucleoid-bound PomZ dimers can diffuse on the nucleoid and, when the dimer is attached to the cluster, also on the PomXY cluster with diffusion constants, $D_{\text{nuc}}$ and $D_{\text{clu}}$, respectively. These two diffusive processes are implemented as stochastic hopping events that occur with rate $k^0_{\text{hop, nuc}} = D_{\text{nuc}}/a^2$ and $k^0_{\text{hop, clu}} = D_{\text{clu}}/a^2$. More concretely, a nucleoid-bound PomZ dimer at site $(i, j)$, with $i$ denoting the lattice site in $x$-direction and $j$ in $y$-direction, can move to site $(i \pm 1, j)$ or $(i, j \pm 1)$ with hopping rate $k^0_{\text{hop, nuc}}$. As we are using periodic boundary conditions in $y$-direction, a particle may also hop from site $(i, N_{\text{nuc},y})$ to the site $(i, 1)$ and vice versa; here $N_{\text{nuc},y} = w_{\text{nuc}}/a$ denotes the number of sites along the $y$-axis. In $x$-direction we assume reflecting boundary conditions for the PomZ dimer movements.

If a PomZ dimer is bound to both cluster and nucleoid, a hopping event leads to gain or loss in elastic energy. In this case we multiply the constant hopping rates by exponential factors, which are chosen such that detailed balance holds (see Lansky et al. [25]):

$$k_{\text{hop,clu}} = k^0_{\text{hop,clu}} \cdot \exp\left[-\frac{1}{4}\beta k \left[(\vec{x}_{\text{clu, new}} - \vec{x}_{\text{nuc}})^2 - (\vec{x}_{\text{clu, old}} - \vec{x}_{\text{nuc}})^2\right]\right] , \tag{S3}$$

$$k_{\text{hop,nuc}} = k^0_{\text{hop,nuc}} \cdot \exp\left[-\frac{1}{4}\beta k \left[(\vec{x}_{\text{clu}} - \vec{x}_{\text{nuc, new}})^2 - (\vec{x}_{\text{clu}} - \vec{x}_{\text{nuc, old}})^2\right]\right] . \tag{S4}$$

The labels "old" and "new" refer to the positions of the binding sites before and after the hopping event.

##### 4. Movement of the PomXY cluster

Cluster-bound PomZ dimers can exert forces, which lead to a net movement of the cluster. Let us denote the position of the nucleoid binding site of the $i$-th PomZ dimer as $\vec{x}_{i,\text{nuc}}$, and the position of the cluster binding site as $\vec{x}_{i,\text{clu}} = \vec{x}_c + \Delta\vec{x}_{i,\text{clu}}$. With $\vec{x}_c = (x_c, y_c) \in \mathbb{R}^2$ we denote the position of the midpoint of the cluster. Here, we decomposed the position of a cluster binding site into two parts: the cluster position, $\vec{x}_c$, and an additional vector $\Delta\vec{x}_{i,\text{clu}}$. The reason for this decomposition is that we are interested in the equation of motion for the cluster position, $\vec{x}_c$, and, as long as the PomZ dimer does not diffuse on the nucleoid or the cluster, the two vectors $\vec{x}_{i,\text{nuc}}$ and $\Delta\vec{x}_{i,\text{clu}}$ are constant.

A single PomZ dimer, bound to the nucleoid and cluster at fixed positions relative to both scaffolds, exerts a force $\vec{F}_i(t)$ on the cluster:

$$\vec{F}_i(t) = -k\left(\vec{x}_{i,\text{clu}}(t) - \vec{x}_{i,\text{nuc}}\right) = -k\left(\vec{x}_c(t) + \Delta\vec{x}_{i,\text{clu}} - \vec{x}_{i,\text{nuc}}\right) . \tag{S5}$$

In a friction dominated regime, the sum over all forces exerted by $N_b$ cluster-bound PomZ dimers has to balance with the friction force acting on the cluster (friction coefficient $\gamma$):

$$\gamma\dot{\vec{x}}_c(t) = \sum_{i=1}^{N_b} \vec{F}_i(t) = -k\sum_{i=1}^{N_b}\left(\vec{x}_c(t) + \Delta\vec{x}_{i,\text{clu}} - \vec{x}_{i,\text{nuc}}\right) . \tag{S6}$$

This equation is solved by separation of variables, yielding

$$\vec{x}_c(t) = \left(\vec{x}_c(t_0) - \vec{x}_f\right)\exp\left(-\frac{N_b k}{\gamma}(t - t_0)\right) + \vec{x}_f , \tag{S7}$$

with

$$\vec{x}_f = \frac{1}{N_b}\left(\sum_{i=1}^{N_b} \vec{x}_{i,\text{nuc}} - \Delta\vec{x}_{i,\text{clu}}\right) . \tag{S8}$$

The initial time is denoted as $t_0$. We find that the cluster approaches the position $\vec{x}_f$, at which no net force is acting on the cluster, exponentially fast with characteristic time $t_{\text{clu}} = \gamma/(N_b k)$.

##### 5. Cytosolic PomZ distribution

To include the cytosolic PomZ dynamics in our model, we reduce the ATPase cycle to three processes: attachment of PomZ-ATP to the nucleoid, detachment of PomZ-ADP at the cluster, and nucleotide exchange. We model the dynamics of PomZ-ATP and PomZ-ADP in the cytosol as one-dimensional reaction-diffusion equations as described in the following. At the position of the cluster, $x_c(t)$, PomZ-ADP is released into the cytosol. Since the PomZ dynamics is a lot faster than the cluster dynamics, we can assume that the cluster is stationary, $x_c(t) = x_c$ on the time scale of the PomZ dynamics. The local increase in cytosolic PomZ-ADP at the cluster position due to detachment facilitated by ATP hydrolysis is approximated as a point source: $s_0\delta(x - x_c)$. The constant $s_0$ depends on the hydrolysis rate $k_h$ and the amount of PomZ dimers bound to the cluster. However, our final result, the normalized steady-state PomZ-ATP distribution will not depend on this constant. In the cytosol, PomZ-ADP exchanges ADP for ATP nucleotides with an effective rate $k_{\text{ne}}$. We assume that cytosolic PomZ in both nucleotide states diffuses with the same diffusion constant, $D_{\text{cyt}}$. However, only the ATP-bound form of PomZ can attach to the nucleoid with a rate $k_{\text{on}}$. In total, we obtain the following coupled partial differential equations for the cytosolic PomZ-ATP ($c_T$) and PomZ-ADP ($c_D$) density:

$$\partial_t c_D(x,t) = D_{\text{cyt}}\partial_x^2 c_D(x,t) - k_{\text{ne}} c_D(x,t) + s_0\delta(x - x_c)\Theta(t) \ , \tag{S9a}$$

$$\partial_t c_T(x,t) = D_{\text{cyt}}\partial_x^2 c_T(x,t) + k_{\text{ne}} c_D(x,t) - k_{\text{on}} c_T(x,t) \ . \tag{S9b}$$

We solved these two differential equations with no-flux boundary conditions for the stationary case. The solution for PomZ-ATP, given the cluster is at position $x_c$, reads:

$$\begin{aligned} c_T(x;x_c) = \tilde{c}_1 \Bigg[ &-\lambda_T \cosh\left(\frac{L_1}{\lambda_T}\right)\cosh\left(\frac{L_2+x}{\lambda_T}\right)\sinh\left(\frac{L}{\lambda_D}\right) + \\ &+ \lambda_D \cosh\left(\frac{L_1}{\lambda_D}\right)\cosh\left(\frac{L_2+x}{\lambda_D}\right)\sinh\left(\frac{L}{\lambda_T}\right)\Bigg] \ , \quad \text{for} \quad -x_c \le x \le 0 \ , \end{aligned} \tag{S10}$$

$$\begin{aligned} c_T(x;x_c) = \tilde{c}_1 \Bigg[ &-\lambda_T \cosh\left(\frac{L_2}{\lambda_T}\right)\cosh\left(\frac{L_1-x}{\lambda_T}\right)\sinh\left(\frac{L}{\lambda_D}\right) + \\ &+ \lambda_D \cosh\left(\frac{L_2}{\lambda_D}\right)\cosh\left(\frac{L_1-x}{\lambda_D}\right)\sinh\left(\frac{L}{\lambda_T}\right)\Bigg] \ , \quad \text{for} \quad 0 \le x \le l_{\text{nuc}} - x_c \ , \end{aligned} \tag{S11}$$

with

$$\tilde{c}_1 = \frac{4 s_0 \lambda_T^2 e^{L(1/\lambda_D + 1/\lambda_T)}}{D_{\text{cyt}}\left(\lambda_D^2 - \lambda_T^2\right)\left(e^{2L/\lambda_D} - 1\right)\left(e^{2L/\lambda_T} - 1\right)} \ . \tag{S12}$$

We chose the coordinate system such that the cluster position, $x_c \in [0, l_{\text{nuc}}]$, is shifted to the origin. The lengths of the cluster-to-nucleoid end distances left and right of the cluster are given by $x_c$ and $l_{\text{nuc}} - x_c$, respectively. Furthermore, we defined the diffusive length scales for PomZ-ADP until it exchanges its ADP for ATP, and PomZ-ATP until it attaches to the nucleoid as $\lambda_D$ and $\lambda_T$, respectively:

$$\lambda_D = \sqrt{\frac{D_{\text{cyt}}}{k_{\text{ne}}}} \quad \text{and} \quad \lambda_T = \sqrt{\frac{D_{\text{cyt}}}{k_{\text{on}}}} \ . \tag{S13}$$

The above solution for the cytosolic PomZ-ATP density holds true for $\lambda_T \neq \lambda_D$. If the two length scales are equal,

$\tilde{c}_1$ becomes singular and hence this case needs to be considered separately. For $\lambda_D = \lambda_T \equiv \lambda$ the solution is given by:

$$
\begin{aligned}
c_T(x;x_c) = \tilde{c}_2 \Bigg[ &(2L_1 - x)\cosh\left(\frac{2L_2+x}{\lambda}\right) - x\cosh\left(\frac{2L+x}{\lambda}\right) + \\
&+ (2L+x)\cosh\left(\frac{x}{\lambda}\right) + (2L_2+x)\cosh\left(\frac{2L_1-x}{\lambda}\right) + \\
&+ 4\lambda\cosh\left(\frac{L_1}{\lambda}\right)\cosh\left(\frac{L_2+x}{\lambda}\right)\sinh\left(\frac{L}{\lambda}\right)\Bigg] \,, \qquad \text{for} \quad -x_c \le x \le 0 \,,
\end{aligned} \tag{S14}
$$

$$
\begin{aligned}
c_T(x;x_c) = \tilde{c}_2 \Bigg[ &(2L_2 + x)\cosh\left(\frac{2L_1-x}{\lambda}\right) + x\cosh\left(\frac{2L-x}{\lambda}\right) + \\
&+ (2L-x)\cosh\left(\frac{x}{\lambda}\right) + (2L_1-x)\cosh\left(\frac{2L_2+x}{\lambda}\right) + \\
&+ 4\lambda\cosh\left(\frac{L_2}{\lambda}\right)\cosh\left(\frac{L_1-x}{\lambda}\right)\sinh\left(\frac{L}{\lambda}\right)\Bigg] \,, \qquad \text{for} \quad 0 \le x \le l_{\text{nuc}} - x_c \,,
\end{aligned} \tag{S15}
$$

with

$$
\tilde{c}_2 = \frac{s_0}{8D_{\text{cyt}}\sinh^2(L/\lambda)} \,. \tag{S16}
$$

With the analytical solution for the PomZ-ATP density in the cytosol, we can now define the attachment rate of a PomZ dimer to the nucleoid, accounting for the cytosolic PomZ distribution. We normalize the steady-state solution for $c_T(x;x_c)$ to obtain a probability density:

$$
p_T(x;x_c) = \frac{k_{\text{on}}}{s_0} c_T(x;x_c) \,. \tag{S17}
$$

The rate for a PomZ dimer to attach to position $(x,y)$ on the nucleoid is then defined by:

$$
k_{\text{on}}(x,y;x_c) \equiv k_{\text{on}}\, p_T(x;x_c) p_T(y) = \frac{k_{\text{on}}}{w_{\text{nuc}}}\, p_T(x;x_c) \,, \tag{S18}
$$

with a uniform distribution, $p_T(y)$, along the short cell axis direction. Approximately, the attachment rate per lattice site is then given by this value multiplied with the lattice spacing $a$ squared.

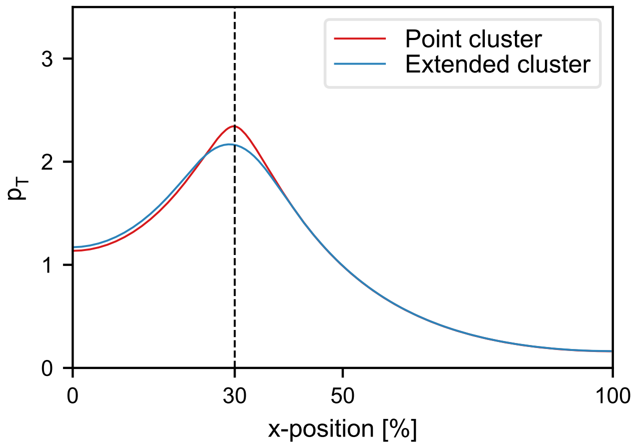


FIG. S1. **Cytosolic PomZ-ATP distribution.** Comparison of the steady-state solutions for the cytosolic PomZ-ATP density if release of PomZ-ADP into the cytosol at the PomXY cluster is modelled as a point source (red line) or a source with the same extension as the cluster (blue line). The cluster is at position $x_c = 30\%$ of nucleoid length. We used the parameters shown in Table S1.

In the presented derivation of the cytosolic PomZ-ATP density we reduced the cytosol to a one-dimensional line and the PomXY cluster to a point source. To investigate how the PomZ-ATP density changes when the cluster's extension is accounted for, we solved equations S9a and S9b with the Dirac delta distribution replaced by a Heaviside step function $\Theta(x - x_c + l_{\mathrm{clu}}/2)\Theta(x_c + l_{\mathrm{clu}}/2 - x)/l_{\mathrm{clu}}$, numerically. We find that the steady-state solution for the PomZ-ATP density, when the cluster is included as a point source (Equations S10 and S11), is a good approximation to the solution considering a one-dimensional cytosolic lane and an extended cluster (Figure S1). The PomZ density profiles only deviate significantly in close proximity to the cluster, which can be attributed to the different shapes of the cluster used.

#### II. DISCUSSION OF PARAMETERS USED IN THE SIMULATIONS

| **Parameter** | **Variable** | **1D model** | **3D model** |
|---|---|---|---|
| Nucleoid length | $l_{\mathrm{nuc}}$ | 5.0 µm | 5.0 µm |
| Nucleoid width (circumference) | $w_{\mathrm{nuc}}$ | - | 2.2 µm |
| PomXY cluster length | $l_{\mathrm{clu}}$ | 0.7 µm | 0.7 µm |
| PomXY cluster width | $w_{\mathrm{clu}}$ | - | 0.7 µm |
| Effective spring stiffness of a PomZ dimer | $k$ | $10^4\,\mathrm{k_B T \mu m^{-2}}$ | $10^4\,\mathrm{k_B T \mu m^{-2}}$ |
| Attachment rate of cytosolic PomZ to nucleoid | $k_{\mathrm{on}}$ | $0.1\,\mathrm{s^{-1}}$ | $0.1\,\mathrm{s^{-1}}$ |
| Attachment rate of nucleoid-bound PomZ to cluster (unstretched) | $k_a^0$ | $500\,\mathrm{s^{-1}\mu m^{-1}}$ | $2.0 \times 10^4\,\mathrm{s^{-1}\mu m^{-2}}$ |
| Diffusion constant of PomZ on nucleoid | $D_{\mathrm{nuc}}$ | $0.1\,\mathrm{\mu m^2 s^{-1}}$ | $0.1\,\mathrm{\mu m^2 s^{-1}}$ |
| Diffusion constant of PomZ on cluster | $D_{\mathrm{clu}}$ | $0.1\,\mathrm{\mu m^2 s^{-1}}$ | $0.1\,\mathrm{\mu m^2 s^{-1}}$ |
| ATP hydrolysis rate of PomZ at the cluster | $k_h$ | $1\,\mathrm{s^{-1}}$ | $1\,\mathrm{s^{-1}}$ |
| Total number of PomZ dimers in the cell | $N$ | 100 | 100 |
| Diffusion constant of the PomXY cluster in the cytosol | $D_{\mathrm{cluster}}$ | $4 \times 10^{-4}\,\mathrm{\mu m^2/s}$ | $4 \times 10^{-4}\,\mathrm{\mu m^2/s}$ |
| Diffusion constant of PomZ in the cytosol | $D_{\mathrm{cyt}}$ | - | $0.1\,\mathrm{\mu m^2 s^{-1}}$, $0.5\,\mathrm{\mu m^2 s^{-1}}$ |
| Nucleotide exchange rate of PomZ | $k_{\mathrm{ne}}$ | - | $6\,\mathrm{s^{-1}}$ |
| Lattice spacing | $a$ | 0.01 µm | 0.01 µm |

TABLE S1. **Parameters used in the simulations.** The parameters used in the one-dimensional model are the same as in [10]. In the three-dimensional model we used the same parameter values when possible.

The total PomZ dimer number, the length and width of the cluster and the length of the nucleoid are chosen in accordance with experimental observations in *M. xanthus* [4]. For the ATP hydrolysis rate we use a value of $1\,\mathrm{s^{-1}}$ as suggested by FRAP experiments where PomZ in the cluster is bleached [4]. The attachment rate of cytosolic PomZ to the nucleoid is approximated by literature values for the related Par system for chromosome and plasmid segregation ($50\,\mathrm{s^{-1}}$ [13] and $0.03\,\mathrm{s^{-1}}$ [12]). To get a double attachment rate that is comparable to the one-dimensional case, we chose $k_a^0$ such that the total attachment rate $k_a^{\mathrm{tot}}$ is equal in the one-dimensional model studied previously [10] and the three-dimensional model studied here. This implies that the rate used in the one-dimensional model has to be multiplied by a factor of $\sqrt{\beta k/2\pi}$ to account for the additional dimension.

The diffusion constant of PomZ on the nucleoid and on the PomXY cluster is approximated by the effective diffusion constant of ParA dimers on the nucleoid used in models for the Par system. However, these values vary a lot: $0.001\,\mathrm{\mu m^2/s} - 1\,\mathrm{\mu m^2/s}$ [13, 26]. The friction coefficient of the cluster, $\gamma$, is related to its diffusion constant, $D_{\mathrm{cluster}}$, via Stokes-Einstein: $\gamma = k_B T/D_{\mathrm{cluster}}$. We approximate the diffusion constant $D_{\mathrm{cluster}}$ by the corresponding literature values for plasmids. However, since the size of the PomXY cluster is larger than the typical size of a single plasmid, these values are only an upper bound for the diffusion constant of the Pom cluster: $D_{\mathrm{cluster}} \leq 10^{-3}\,\mathrm{\mu m^2 s^{-1}}$[13, 14]. The effective spring stiffness of the PomZ dimers, $k$, we approximate by the stiffness of a bond between a plasmid and the nucleoid via ParA dimers [26].

We approximate the nucleotide exchange rate of PomZ-ADP to PomZ-ATP by the corresponding rate for MinD proteins, $6\,\mathrm{s^{-1}}$ [15]. Based on the large diffusion constant of Min proteins in the cytosol (on the order of $10\,\mathrm{\mu m^2 s^{-1}}$, [15]), and the fast dynamics of PomZ in the cytosol as observed in FRAP experiments [4], we expect the cytosolic diffusion constant of PomZ to be large. When we tested the effect of a non-homogeneous PomZ-ATP distribution in the cytosol on the cluster dynamics, we chose diffusion constants that are two orders of magnitudes smaller ($0.1\,\mathrm{\mu m^2/s}$ or $0.5\,\mathrm{\mu m^2/s}$).

#### III. DETAILS ON THE STOCHASTIC SIMULATION

##### 1. Initial PomZ distribution

Initially, all PomZ proteins are in the cytosol. Then we let the simulations run for a time $t_{\min}$ of at least 10 minutes with a fixed cluster position such that the PomZ proteins can approach their steady-state distribution. The time $t_{\min}$ is chosen such that it is larger than the typical time scales for the PomZ dynamics, i.e. the time scale for attachment to the nucleoid ($1/k_{\text{on}} = 10\,\text{s}$), the time scale for PomZ to explore the whole nucleoid by diffusion ($l_{\text{nuc}}^2/2D_{\text{nuc}} = 125\,\text{s}$), and the time scale for cluster-bound PomZ to detach ($1/k_h = 1\,\text{s}$). After the initial time, $t_{\min}$, recording starts. The cluster can now start to move or is kept at a fixed position ("stationary simulation") during the entire simulation.

##### 2. Gillespie algorithm

We implemented our model using the Gillespie algorithm [10, 27], a stochastic simulation algorithm. Since the cluster position, $\vec{x}_c \in \mathbb{R}^2$, changes over time according to the forces cluster-bound PomZ dimers exert on the cluster (Equation S7), all rates that depend on the position of the cluster binding site of a PomZ dimer, depend on time. These include the attachment rate of a nucleoid-bound PomZ dimer to the cluster and the hopping rates of a PomZ dimer bound to the nucleoid and the cluster. However, if the cluster only moves slightly in one time step of the Gillespie algorithm, we can approximate the time-dependent rates as constant.

To quantify the effect of the time dependence of the rates, let us consider a PomXY cluster with $N_b$ PomZ dimers bound to it such that a non-zero net force acts on the cluster. According to Equation S7 the time scale for the cluster to relax to the force-free position, $\vec{x}_f$, is given by $t_{\text{clu}} = \gamma/(N_b k)$. The number of cluster-bound PomZ dimers, $N_b$, changes with the position of the cluster along the nucleoid. For a cluster positioned at 10% of nucleoid length and one at mid-nucleoid the number of cluster bound PomZ dimers, as obtained from simulations, leads to $t_{\text{clu}} \approx 0.18\,\text{s}$ and $t_{\text{clu}} \approx 0.09\,\text{s}$, respectively (for the parameters as in Table S1).

Next, we consider the time step, $\Delta t$, until the next event happens in the Gillespie algorithm. The most frequent event is hopping of PomZ dimers on the nucleoid for the parameters we consider (Table S1). Hence, the time step $\Delta t$ can be approximated by the time until a PomZ dimer bound to the nucleoid, hops on the nucleoid. The rate for the event that any of the nucleoid-bound PomZ dimers, $N_{\text{nuc}}$, hops in any of the four possible directions on the nucleoid (ignoring the boundaries) is given by $4\,k_{\text{hop,nuc}}^0 N_{\text{nuc}}$. The typical time until the next event happens can then be approximated by the inverse of this rate. Again, the number of nucleoid-bound PomZ dimers, $N_{\text{nuc}}$, varies with the position of the cluster. For a cluster at 10% of nucleoid length and one at mid-nucleoid, we get $\Delta t \approx 3 \times 10^{-6}\,\text{s}$ and $\Delta t \approx 4 \times 10^{-6}\,\text{s}$, respectively. Since the typical time until the next event happens, $\Delta t$, is much smaller than the time scale for the movement of the cluster, $t_{\text{clu}}$, we can approximate all rates in the Gillespie algorithm as time-independent, which significantly improves the computational speed of the algorithm.

#### IV. PROCESSING OF SIMULATED DATA

##### 1. PomZ flux on the nucleoid

The PomZ flux along the nucleoid for a specific cluster position is determined by recording the PomZ flux at any time the cluster is in a small region ($\pm 0.5\%$ of nucleoid length) around the $x$-position of the cluster of interest. To obtain the flux of PomZ into the cluster along the long cell axis direction, the fluxes are averaged over the values in y-direction, but only considering the region of the nucleoid that corresponds to the extension of the cluster region along the long cell axis (yellow region in Figure S2A). Additionally, the data is averaged over an ensemble of about 100 simulations. An example for such an averaged flux profile is shown in Figure S2B. To obtain the difference in the PomZ fluxes into the cluster from each side along the long cell axis, the maximal / minimal values of the average flux profile left / right of the PomXY cluster are determined (red lines in Figure S2B). Finally, the two flux values of different signs are added together to obtain the flux difference.

#### V. FLUX DIFFERENCE INTO THE CLUSTER

PomZ dimers detach from the nucleoid into the cytosol upon ATP hydrolysis, which breaks detailed balance and leads to a net flux of PomZ in the system. In the following we consider the fluxes of PomZ for the one- and three-

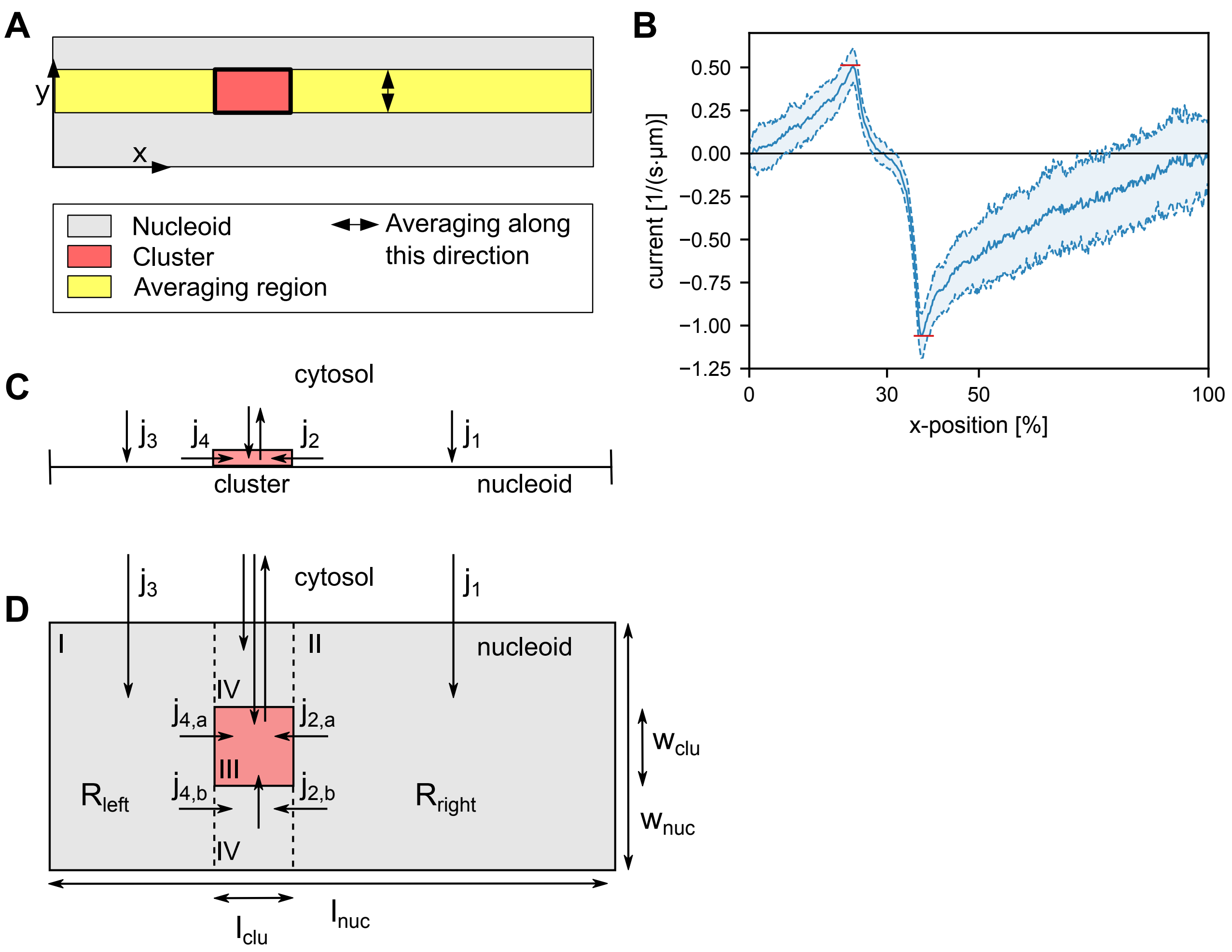


FIG. S2. **Flux of PomZ in the three-dimensional model geometry.** (A) Sketch of the region used for averaging the flux. Only hopping events in the yellow region are taken into account when determining the PomZ flux profile along the long cell axis. The fluxes are averaged over the values in $y$-direction. (B) Simulated flux profile of nucleoid-bound PomZ along the long cell axis with averaging performed as illustrated in (A). The flux into the cluster is given by the maximal value left, and the minimal value right of the cluster (red horizontal lines). We used the same parameters as in Table S1. (C) In the one-dimensional model, nucleoid and cluster are incorporated as one-dimensional lattices. The cluster region is shown as a red rectangle. PomZ dimers attach to and diffuse on the nucleoid with reflecting boundary conditions at the nucleoid ends. The arrows show the different PomZ fluxes from one region (cytosol, cluster and nucleoid left and right of the cluster) to another. (D) Similar to (C), but the fluxes in the three-dimensional model geometry are shown. The grey region shows the nucleoid of size $l_{\text{nuc}} \times w_{\text{nuc}}$ and the red region the cluster of size $l_{\text{clu}} \times w_{\text{clu}}$. The area of the nucleoid regions left and right of the cluster are denoted by $R_{\text{left}}$ and $R_{\text{right}}$, respectively.

dimensional model geometry (see Figure S2C,D). Since the cluster is a lot less mobile than the PomZ dimers, we can approximate the cluster position as stationary. In this case, the fluxes in and out of each region (cytosol, cluster and nucleoid region left and right of the cluster) have to balance in the steady state. In the one-dimensional geometry, the fluxes of PomZ dimers to the nucleoid right and left of the cluster, $j_1$ and $j_3$, balance the fluxes into the cluster region, $j_2$ and $j_4$, respectively (Figure S2C). If PomZ is homogeneously distributed in the cytosol, the fluxes onto the nucleoid scale with the lengths of the respective nucleoid regions:

$$j_1 = k_{\text{on}} \frac{l_{\text{nuc}} - x_c - l_{\text{clu}}/2}{l_{\text{nuc}}} N_{\text{cyt}} \ , \tag{S19}$$

$$j_3 = k_{\text{on}} \frac{x_c - l_{\text{clu}}/2}{l_{\text{nuc}}} N_{\text{cyt}} \ , \tag{S20}$$

with $x_c$ the position of the cluster, and $N_{\mathrm{cyt}}$ the number of PomZ dimers in the cytosol. This results in the following formula for the flux difference of PomZ into the cluster

$$j_{\mathrm{diff}} = j_2 - j_4 = j_1 - j_3 = k_{\mathrm{on}} N_{\mathrm{cyt}} \left(1 - \frac{2x_c}{l_{\mathrm{nuc}}}\right) . \tag{S21}$$

The flux difference is proportional to the attachment rate of PomZ to the nucleoid, $k_{\mathrm{on}}$, and the number of PomZ dimers in the cytosol, $N_{\mathrm{cyt}}$. It is important to note, that $N_{\mathrm{cyt}}$ also depends, among other parameters, on the position of the cluster $x_c$.

In the three-dimensional model geometry there are additional fluxes compared to the one-dimensional geometry if the cluster does not encompass the entire nucleoid circumference, i.e. if $w_{\mathrm{clu}} < w_{\mathrm{nuc}}$ holds (see Figure S2D). Nucleoid-bound PomZ dimers in region I or II can leave these regions either by entering the cluster region and then attaching to the cluster, or by diffusing into the region in the extension of the cluster along the short cell axis (region IV in Figure S2D). In the latter case, the PomZ dimers can enter the cluster region along the short axis, diffuse back into the region they came from or diffuse past the cluster. In the steady state, the fluxes in and out of each region have to balance. For region I and II this implies:

$$j_1 = j_{2,a} + j_{2,b} \ , \tag{S22}$$
$$j_3 = j_{4,a} + j_{4,b} \ . \tag{S23}$$

If we assume that the fluxes into the cluster region (region III) and into region IV scale with the extensions of the respective regions along the short cell axis, i.e. $j_{2,a}/j_{2,b} = w_{\mathrm{clu}}/(w_{\mathrm{nuc}} - w_{\mathrm{clu}})$ and similarly for $j_{4,a}$ and $j_{4,b}$, the flux difference into the cluster reads:

$$j_{\mathrm{diff}} = j_{2,a} - j_{4,a} = k_{\mathrm{on}} N_{\mathrm{cyt}} \frac{w_{\mathrm{clu}}}{w_{\mathrm{nuc}}} \left(1 - \frac{2x_c}{l_{\mathrm{nuc}}}\right) \ , \tag{S24}$$

which agrees with the formula for the one-dimensional system if the cluster is ring-shaped, i.e. $w_{\mathrm{clu}} = w_{\mathrm{nuc}}$. This analytical expression fits well with our simulation results for a ring-shaped cluster (see Figure 3D). However, if the cluster does not cover the full nucleoid's width, it deviates from the simulation results. This deviation can be attributed to the fact that PomZ dimers that diffuse into region IV are not absorbed here, but can diffuse back into regions I or II. Hence, the fluxes into the cluster are larger than the values obtained from the simple estimate we used before. For the fluxes into the cluster from the right, we have

$$\frac{j_{2,a}}{j_{2,b}} > \frac{w_{\mathrm{clu}}}{w_{\mathrm{nuc}} - w_{\mathrm{clu}}} \ . \tag{S25}$$

Furthermore, PomZ dimers can pass the cluster by diffusion from region I to II or vice versa. However, this flux only matters if the cluster is small both in length and width.

#### VI. CLUSTER MOVEMENT ALONG SHORT CELL AXIS DIRECTION

In addition to the cluster dynamics along the long cell axis, as discussed in the main text, we also considered the dynamics along the short cell axis. Because of the rotational symmetry the clusters do not have a preferred direction along this axis, on average. However, both diffusive and persistent, unidirectional motion is conceivable. We expect persistent movement if PomZ diffusion on the nucleoid is slow compared to the cluster dynamics, such that there is a delay between the cluster movement and the PomZ gradient reaching its steady state [10, 14, 28 - 30]. In this case, the initial direction of the cluster's movement along the nucleoid's circumference is chosen stochastically by the interactions of the PomZ dimers with the cluster. Once the cluster started to move in one direction, it is more likely to continue in this direction, because the asymmetry in the PomZ density gradient is maintained. In contrast, for fast PomZ diffusion on the nucleoid, we expect an approximately symmetric PomZ distribution around the cluster, resulting in equal likelihood for the cluster to move in each direction, suggesting diffusive motion. For the parameter set we considered (Table S1) the clusters show diffusive behavior in $y$-direction as indicated by a mean-square displacement that grows linearly in time (Figure S3, inset).

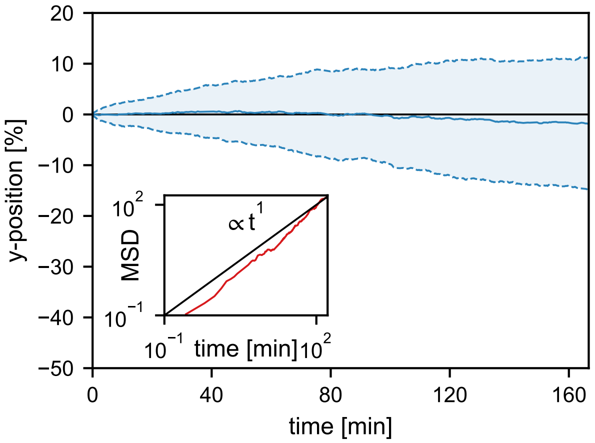


FIG. S3. **Cluster movement along short cell axis direction.** Average cluster trajectory in $y$-direction using an ensemble of 100 simulations (solid blue line). The shaded region indicates one standard deviation above and below the average trajectory. The inset shows the mean-square displacement, which increases linearly in time, indicating diffusive motion.

#### VII. DYNAMICS OF TWO POM CLUSTERS

Motivated by equidistant positioning of plasmids by ParAB$S$ systems we investigated the dynamics of two Pom clusters in the realistic three-dimensional cell geometry. We find that two clusters localize at the one- and three-quarter positions along the nucleoid, i.e. at equidistant positions (Figure S4A).

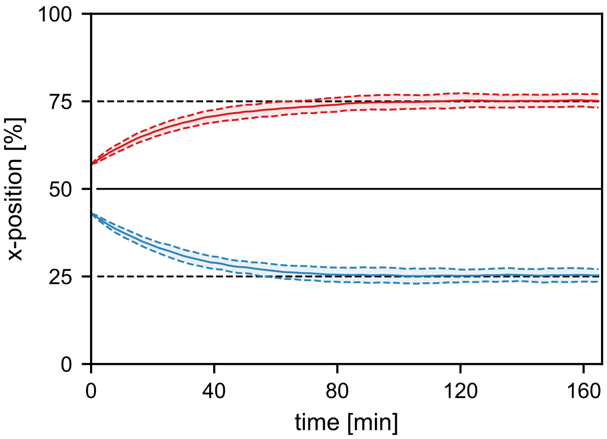


FIG. S4. **Two clusters are localized at the one- and three-quarter positions.** Averaged positions of the two clusters along the $x$-direction (solid lines, ensemble of 100 runs). The shaded regions indicate the average cluster positions plus and minus one standard deviation. Initially, the two clusters are positioned side by side along the long cell axis such that the left edge of one and the right edge of the other cluster are positioned at mid-nucleoid (see also Movie S2). We used the parameter set given in Table S1.

Flux-based positioning of two clusters can be heuristically explained as follows (see also [13, 14, 21]): If two protein clusters approach each other, the PomZ density and fluxes on the nucleoid between the clusters are reduced. Since the clusters move into the direction of the highest PomZ flux, two clusters effectively repel each other, which separates the clusters from each other. Moreover, the average positions of the two clusters and the average PomZ density on the nucleoid has to be symmetric with respect to the mid-nucleoid plane. Hence there is no net flux of nucleoid-bound PomZ at mid-nucleoid, on average. Therefore, in the stationary state, we can map the system to one that consists of two subsystems with half the size of the original system and each subsystem contains one cluster. The positioning mechanism previously discussed for one cluster explains midcell localization in each of the subsystems, which corresponds to the one- and three-quarter positions on the nucleoid.

#### VIII. MOVIES

**Movie S1**: The movie shows the simulated position of one PomXY cluster together with the PomZ density on the nucleoid over time. The black rectangle indicates the contour of the cluster and the color scale shows a high density of PomZ in red, a low density in yellow. The cluster starts at one of the nucleoid ends such that the cluster still fully overlaps with the nucleoid. Initially, all PomZ dimers are in the cytosol and have time to equilibrate while the cluster is at a fixed position for 1000 s. Then, the cluster starts to move and we simulated the cluster and PomZ dynamics for 20 000 s. We used the same parameters as in Table S1.

**Movie S2:** Same as Movie S1, but for a simulation with two clusters and a shorter simulation time of 10 000 s. The two clusters start side-by-side along the long cell axis such that the left edge of one and the right edge of the other cluster coincides with mid-nucleoid. The parameters as given in Table S1 were used.
